# Ly-6A-Induced Growth Inhibition and Cell death in a Transformed CD4^+^ T cell line: Role of Tumor Necrosis Factor-α (TNF-α)

**DOI:** 10.1101/2021.11.16.468830

**Authors:** Akshay G. Patel, Sarah Moxham, Anil K. Bamezai

## Abstract

Engaging the Ly-6A protein, an inhibitory signaling protein, on CD4^+^ T cell lines triggers apoptosis. Signaling through Ly-6A activates cell-intrinsic apoptotic cell death pathway as indicated by release of cytochrome C, activation of caspase 3 and 9. In addition Ly-6A induces cytokine production and growth inhibition. The mechanism underlying simultaneous distinct cellular responses has remained unknown. To examine the relatedness of distinct responses generated by engaging Ly-6A, we have quantified the secretion of TNFα, TGFβ and a related protein GDF10, the three pro-apoptotic, growth inhibitory and tumor suppressive cytokines. While low levels of TGFβ and GDF10 were detected after engaging Ly-6A, the production of TNF-α was elevated in cell cultures stimulated the Ly-6A protein. Blocking the biological activity of TNFα resulted in reduced apoptosis induced by engaging Ly-6A. In contrast, growth inhibition/apoptosis in response to antigen receptor complex stimulation was not observed. Engaging the antigen receptor through activating the epsilon (ε) chain of CD3 generated high levels of TGF-β and GDF10 while decreasing TNFα. These results suggest that the TNF-α cytokine contributes to the Ly-6A-induced growth inhibitory and pro-apoptotic response in CD4^+^ T cells and provides mechanistic explanation of the observed biologically distinct responses initiated after engaging Ly-6A protein. These findings aid in understanding the inhibitory signaling initiated by Ly-6A protein, especially in the context of its potential immune checkpoint inhibitory role in T cells.

## INTRODUCTION

Ly-6A, a member of Ly-6 gene family, is reported to possess cell adhesion and cell signaling properties [1–5]. Based on *in vitro* experiments with primary CD4^+^ T cells, Ly-6A overexpression inhibits antigen receptor-stimulated clonal expansion of primary CD4^+^ T cells [6]. While the primary CD4^+^ T cells overexpressing Ly-6A protein show a hypo-proliferative response to a specific antigen, the CD4^+^ T cells from Ly-6A deficient mice are moderately hyper-responsive to anti-TCR/CD3 stimulation [7]. Engaging the Ly-6A protein on transformed clonal murine CD4^+^ T cell lines with anti-Ly-6A antibody induces distinct cellular processes including activation as assessed by generation of cytokines, apoptosis and growth inhibition [6–10]. The growth inhibited cells show production of the cytokine IL-2 and upregulation of the inhibitory cell cycle protein, p27^kip^ without change in p53 expression [10]. Additionally, Ly-6A signaling induced death in murine T cell lines through a cell-intrinsic apoptotic mechanism as indicated by the translocation of cytochrome C to the cytoplasm and the presence of activated caspases 3 and 9 [10]. We sought to investigate the underlying mechanism for the functionally distinct responses generated in the murine T cell lines. We considered the two possibilities. First, by engaging Ly-6A, multiple distinct signaling pathways were initiated that resulted in varied functional responses (cytokine production, apoptosis etc.,) through a direct cell-autonomous mechanism. Alternatively, Ly-6A induces apoptosis indirectly, through a non-cell autonomous mechanism, first by secreting growth inhibitory/cell death factors that in turn trigger growth inhibition/apoptosis. We report that Ly-6A-induced growth inhibition and apoptosis partly occurs by a non-cell autonomous mechanism mediated by TNF-α generated after engaging the Ly-6A protein on the clonal CD4^+^ T cell line.

## MATERIALS AND METHODS

### Cell lines and antibodies

YH16.33, a CD4^+^ T-cell hybridoma cell line, was used in our assay [11]. Monoclonal antibodies against the epsilon chain of the CD3 protein (145-2C11) [12] and Ly-6A (8G12) [13] were used to stimulate YH16.33. MVB2, a CD4^+^ T cell hybridoma cell line lacking GPI-anchors, and thus lacking Ly-6A on its surface, served as a control [10].

### Cell Culture/Growth Inhibitory Assays

YH16.33 were expanded in RPMI 1640-based cell culture media supplemented with 10% FBS, 2 mM of non-essential amino acids, 100 IU/ml penicillin and 100 IU/ml streptomycin and 0.25 μg/ml Amphotericin B and 2 mM HEPES (Thermo Fisher Scientific, Waltham, MA, USA). Cells were cultured in the presence or absence of antibodies in 5% CO_2_ incubator at 37°C with humidity for 24 hours. Experiments were set with YH16.33 by expending the passaged cells with fresh media one day prior to the day of the experiment. For experimental cultures, 2.5 × 10^4^ cells were cultured in each well of 96-well plate (CytoOne, USA Scientific, Orlando, FL, USA) with or without antibodies and/or other treatments in final volume of 200 μL of culture media.

### Growth Inhibition Assays

MTT assay (Promega, Madison, WI, USA). MTT reagent containing tetrazolium salt 3-(4,5-dimethylthiazol-2-yl)-2,5-diphenyltetrazolium bromide), was used to quantify growth inhibition and proliferation in our cultures. The blue formazan product observed in cell cultures was assessed by spectrophotometer plate reader at 490 nm [14]. The MTT assay quantifies overall metabolic activity that shows a tight association with T cell expansion and response [15,16]. The relative viability in cell cultures was calculated by normalizing the absorbance values obtained in the treatment group to the absorbance of the “No Treatment” group using the following formula:

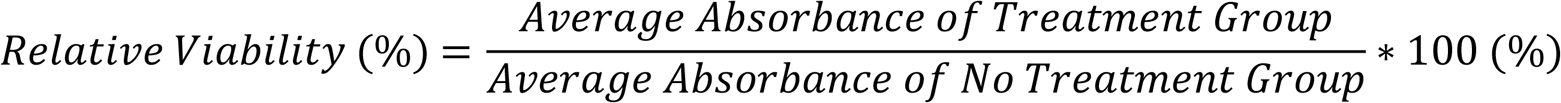

The “No Treatment groups” were assigned a relative viability of 100%, while the TNFα, TGFβ, and GDF10 cytokines in the supernatants from the control and treatment group cell cultures were quantified (see below).

### Cytokine Assays

**S**upernatant (100 μL) from control and treatment cultures were harvested at both 24- and 48-hour time points and mixed with 400μL of culture media in a 96-deep well storage plate (VWR International, Radnor, PA, USA) and stored frozen at −20°C for cytokine assays by ELISA. The cytokine TNFα (BD Bioscience, San Jose, CA, USA), TGFβ (Total and Free Active), (BioLegend, San Diego, CA), and GDF10 (MyBioSource, San Diego, CA, USA) assay kits were purchased from the vendor and cytokines were estimated as per vendor’s instructions. Total TGFβ levels were quantified after acidification, neutralization, and further dilution steps to free TGFβ from its inactivating LAP protein [17] prior to quantification by ELISA.

### Antibody Blocking Assays

To assess the role of the TNFα cytokine in growth inhibition of CD4^+^ T cells induced by the anti-Ly-6A antibody, the biological activity of TNFα was neutralized with an anti-TNFα antibody (Clone MP6-XT22) (BioLegend, San Diego, CA, USA). Cultures with isotype-matched antibody (BioLegend, San Diego, CA, USA) served as controls for determining the specificity in these blocking assays.

### Apoptotic Quantification

Apoptosis in cell cultures was assessed after staining with Annexin V-FITC (BioLegend, San Diego, CA, USA) and Propidium Iodide (PI) (BD Bioscience, San Jose, CA, USA) followed by flow cytometry using FACS Calibur (BD Biosciences, San Jose, CA, USA) as previously published (Lang et.al 2017). For each analysis, 10,000 cells were acquired in FL1 (FITC) and FL2 (PE) channels, respectively, without any gating protocols. The absence of staining with Annexin V-FITC and PI enumerates the live cell population. Cells with bound Annexin V-FITC without PI staining mark cells undergoing apoptosis but the plasma membrane is impervious to PI (early apoptosis). The Annexin V-FITC and PI double positive cells were deemed cells that had undergone apoptosis.

### Statistical Analysis

ANOVA analysis with Tukey-Kramer post-test analysis were used to determine significance in MTT cell proliferation assay and flow cytometric analyses. ANOVA analysis showed us the differences among group means, while the Tukey-Kramer post-test analysis quantified these differences; the null hypothesis was rejected if the p value was <0.05. To determine the significance of cytokine levels from ELISA analysis, two-way ANOVA analyses were performed with a Tukey-Kramer post-test analysis. Connecting letters were reported in conjunction with p-values to show significant differences in the cytokine levels among various antibody treatments (treatment groups sharing connecting letters do not show significant difference, whereas treatment groups with different connecting letters are significantly different).

## RESULTS

### Ly-6A signaling promotes production of TGFβ and GDF10 cytokines in mouse CD4^+^ T Cell Lines

Engaging Ly-6A promotes cytokine production (IL-2 and IFN-γ) and concurrent cell death and growth inhibition in CD4^+^ T cell lines [10]. In order to understand how functionally opposing responses are triggered by Ly-6A signaling in a clonal immortalized CD4^+^ T cell line, we first quantified two growth inhibitory cytokines, TGF-β (as biologically inactive dimer and active monomer) (Figure 1A & B), and GDF10, a member of TGF-β family (Figure 1C). An average of 8.8±4.2 and 11±6.7 ng/mL total TGFβ was detected at 24- and 48-hour time points, respectively. TGFβ produced in response to Ly-6A was not significantly different from than the untreated cultures that served as background controls (7.8±3.1 ng/mL at 24 hours, p=0.9989; and 8.5±5.6 ng/mL, p=0.9306 at 48 hours). In contrast, anti-CD3ε generated an average of 37.5±9.2 ng/mL and 36.3±10.9 ng/mL of total TGFβ at 24 and 48 hours respectively, over 3.5-fold higher response than the untreated controls. The amount of TGFβ detected in anti-CD3ε treated cultures was significantly higher (p<0.0001) than the untreated controls and anti-Ly-6A treated cultures (p<0.0001) (Figure 1A). We quantified the free active monomeric form of TGFβ in the same cell cultures. An average of 59±5 pg/mL and 52±5 pg/mL of free active TGFβ was detected in cell cultures treated with anti-Ly-6A antibody at 24- and 48-hour time points, respectively. The active TGF-β generated in anti-Ly-6A cultures was not significantly different from the untreated cell cultures over the same time course (Figure 1B). Anti-CD3ε treated cell cultures generated 106±47 and 102±65 pg/mL of active TGFβ at 24- and 48-hour time points, respectively, which was significantly higher than the untreated controls (24 hour: p=0.0466; 48 hours: p=0.0134) (Figure 1B).

**Figure 1.**
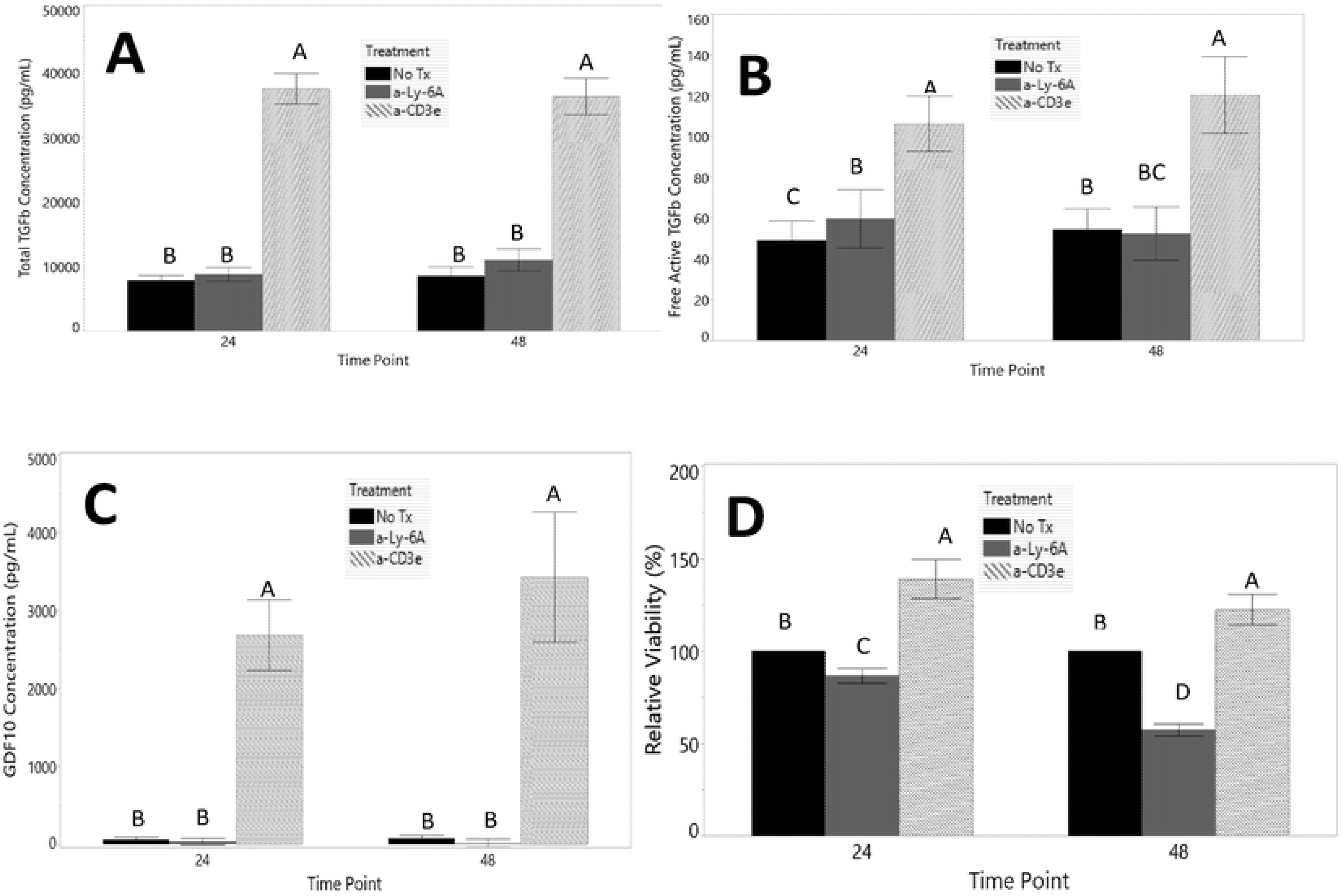
Production of Total TGFβ, free (active) TGFβ and GDF-10 by YH16.33 cells in response to activation through Ly-6A and CD3ε. YH16.33 cells were treated with anti-Ly-6A (gray), anti-CD3ε (white), or left untreated (black) (control) for 24 and 48 hours, and supernatants were harvested for total TGFβ (A), free TGFβ (B) and GDF-10 (C) quantification by ELISA. Growth inhibitory response of YH16.33 cells in the same assay is shown (D). Data are indicative of triplicates run during 4 (active TGFβ) or 5 (total TGFβ and GDF-10) independent trials. The mean of each treatment group is plotted, with error bars showing standard error. ANOVA analysis was performed with a post-test Tukey test. Groups with dissimilar connecting letters are significantly different from each other (p<0.05); n= 4 or 5.

We next sought to assay GDF10, a TGF-β-like cytokine and another reported ligand for TGF-βR1/II [18]. Ly-6A expression is reported to promote tumorigenesis in epithelial cells by influencing signaling through GDF-10 and TGFβRI/II receptor signaling axis [19–21]. An average of 37±165 and 16±204 pg/mL of GDF10 was detected in cell cultures treated with anti-Ly-6A at 24- and 48-hours post-treatment, respectively (Figure 1C). The amount of GDF10 present in anti-Ly-6A treated cultures was not significantly different from the untreated controls (24-hour: 60±1 pg/mL and 48-hour: 75±1 pg/mL). In contrast, 45 – 70-fold higher GD-F10 (24-hour: 2684±1757 and 48-hour: 3429±3236 pg/mL) was detected in anti-CD3ε cultures than the untreated controls during the same time course (Figure 1C).

In summary, our data show that in response to Ly-6A signaling, YH16.33 cells produce insignificant amounts of GDF10 and TGF-β, two known growth inhibitory cytokines that are equivalent to the untreated controls, and therefore less likely to be involved in the observed Ly-6A-mediated growth inhibition (Lang et.al., Figure 1D). This inference is consistent with the observation that significantly higher amounts of GDF10 and TGFβ are detected in YH16.33 cell cultures in the presence of anti-CD3ε, where growth inhibition was not observed (Figure 1D).

### Engaging Ly-6A Expressed on YH16.33 with anti-Ly-6A antibody induces production of TNFα

TNF-α, along with other members of TNF gene family, are known to have, depending on the context of stimulation, either growth inhibitory/pro-apoptotic or cell survival effects [22,23]. To examine the role of TNFα in Ly-6A-induced growth inhibition in YH16.33 cells, we quantified TNFα in the Ly-6A-dependent growth-inhibited cell cultures (Figure 2). An average of 469±2 pg/mL and 594±2 pg/mL of TNFα was detected at 24- and 48-hour time points, respectively (Figure 2A). TNF-α generated in cell cultures with anti-Ly-6A was significantly (p<0.0001) higher than the control untreated cell cultures (24 hours: 60±41 pg/mL; 48 hours:184±344 pg/mL). An average of 217±5 pg/mL and 395±1 pg/mL was detected at 24- and 48-hour time points, respectively, in cultures with anti-CD3ε antibody. Anti-CD3ε induced TNFα levels were higher than the untreated cell cultures; however, it was significantly lower than amounts generated in the presence of Ly-6A antibodies (24 hours: p=0.0033; 48 hours: p=0.0366) (Figure 2A).

**Figure 1.**
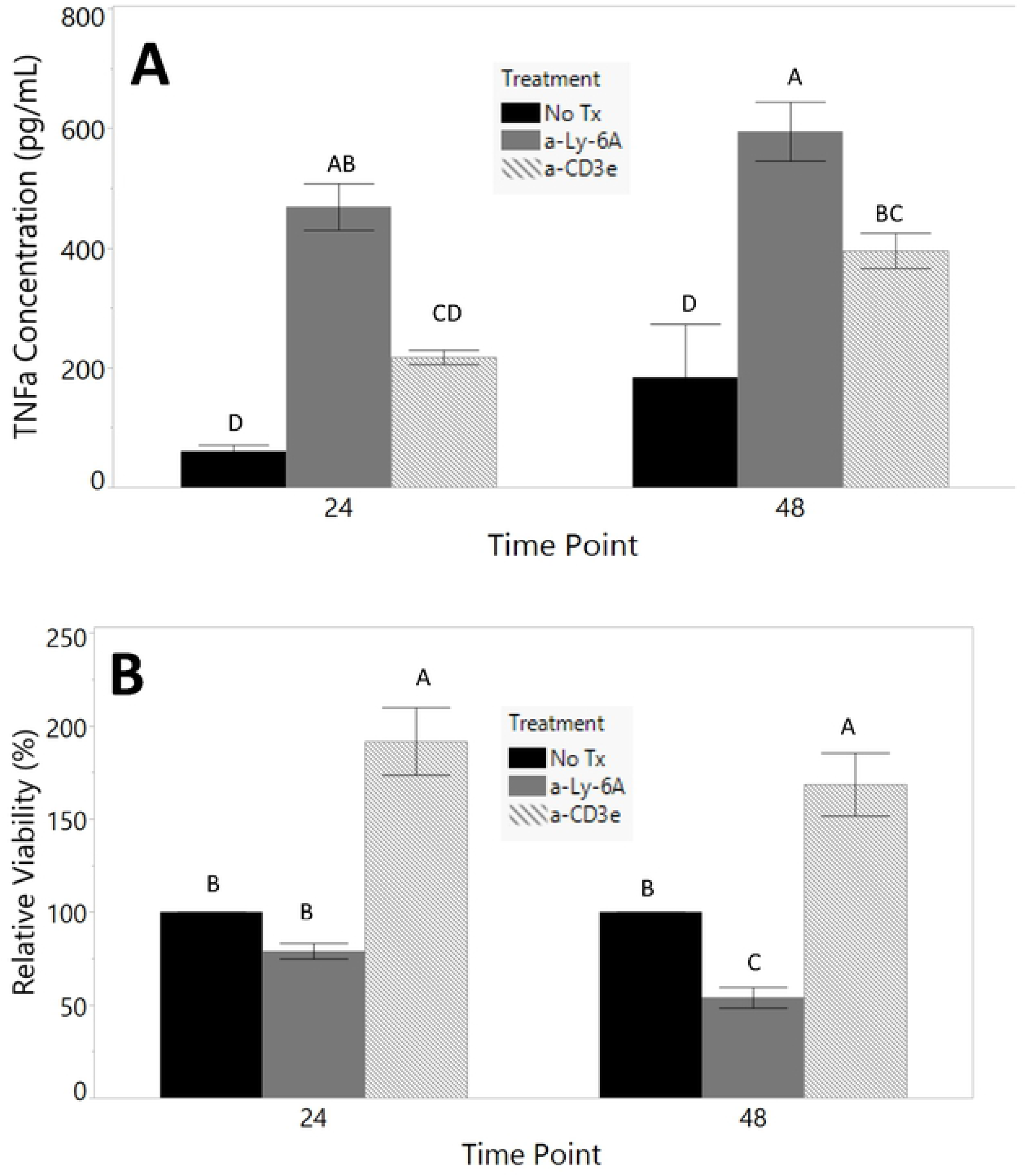
Engaging Ly-6A induces Production of TNFα in YH16.33 cells. YH16.33 cells were treated with anti-Ly-6A (gray), anti-CD3ε (white), or left untreated (black) (control) for 24 and 48 hours, and supernatants were harvested for TNFα quantification by ELISA (A) and growth inhibition using MTT assay (B). Data are indicative of triplicates run during 5 independent trials. The mean of each treatment group is plotted, with error bars showing standard error. ANOVA analysis was performed with a post-test Tukey test. Groups with dissimilar connecting letters are significantly different from each other (p<0.05); n=5.

### Neutralizing biological activity of TNFα partially restores Ly-6A triggered growth-inhibition in YH16.33 cells

We next sought to examine the functional role of TNFα in anti-Ly-6A induced growth inhibition and apoptosis by including neutralizing anti-TNFα antibody in YH16.33 cell cultures. The relative viability of YH16.33 cells treated with the anti-Ly-6A antibody in the absence of anti-TNF-α antibody was significantly reduced (Figure 3), as expected [10]. Inclusion of a biologically active anti-TNFα monoclonal antibody (mAb) in these cultures resulted in moderately enhanced cell survival in anti-Ly-6A treated cell cultures when assessed at 24-hour time point However, this increase was not significant. At the 48-hour time point, a 72% increase in viable cells was observed in cultures with 30μg/mL anti-TNF-α antibody than the untreated control group (p=0.0049). Moderately higher cell viability was observed at the lower concentration of anti-TNF-α (3μg/mL), but this was not statistically significant. Though treatment with anti-TNFα neutralizing antibody at 30μg/mL concentration at 48 hours increased the average relative viability of cells in anti-Ly-6A stimulated cell cultures, the viability was still significantly less than that of the untreated group (p=0.0003) (Figure 3). Taken together, our data indicates that the inclusion of a higher concentration of anti-TNFα antibody significantly reversed the Ly-6A-induced growth inhibition in YH16.33 cells at 48-hour cell cultures.

**Figure 3.**
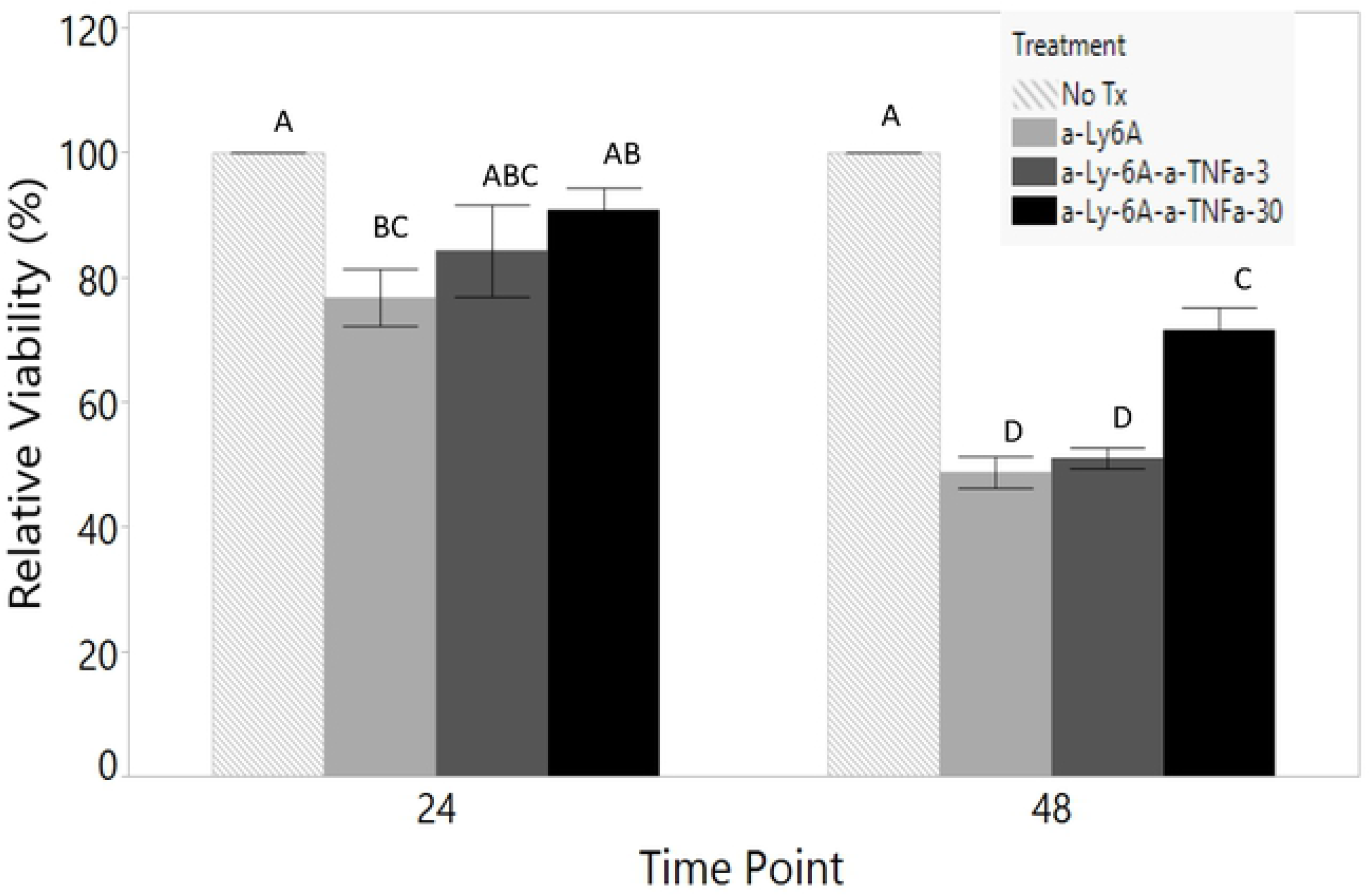
Effect of neutralizing anti-TNFα antibody on viability of anti-Ly-6A stimulated YH16.33 cells. Relative Viability of YH16.33 in response to treatment with anti-Ly-6A (light gray bar), anti-Ly-6A with TNFα neutralizing antibody at 3μg/mL (dark gray bar), anti-Ly-6A with TNFα neutralizing antibody at 30μg/mL (black bar) and untreated cells (white bar) at 24 and 48 hours. Cells were treated as specified and proliferation was assessed via MTT assay. The above data are relative to the no treatment control group proliferation and are indicative of triplicates run during 4 independent trials. The mean of each treatment group is plotted, with error bars showing standard error. ANOVA analysis was performed with a post-test Tukey test. Groups with dissimilar connecting letters are significantly different from each other (p<0.05); n=4

### Neutralizing anti-TNFα monoclonal antibody partially restores Ly-6A induced growth inhibition by reducing cell death

To assess apoptotic cell death in cell cultures, we stained cells with Annexin V-FITC and Propidium Iodide (PI) followed by flow cytometric analyses. About 27% of the total anti-Ly-6A treated cells were viable (AnnV^neg^, PI^neg^), this fraction was significantly decreased (p=0.0009) when compared to the untreated control cultures to the untreated controls (Figure 4A). Consistent with this result is the finding that the remainder 73% (average) of the Ly-6A antibody treated YH16.33 cell cultures showed apoptotic (Annexin V^pos^ PI^neg^) or dead (Annexin V^pos^ PI^pos^) phenotype at the 48-hour time point (Figure 4B). The combined fraction of dead cells, including the apoptotic (Annexin V^pos^PI^neg^) cells in anti-Ly-6A antibody-treated cultures was significantly higher than the untreated control, as previously reported [10]. Adding neutralizing anti-TNFα antibody, at 3μg/mL and 30μg/mL, to these cultures increased the proportion of viable cells to 41% and 43% respectively (Figure 4A). However, this increase was not significant when compared to the anti-Ly-6A treated cultures alone or in combination with isotype control antibody (Figure 4A). An average of 58% and 57% cells in these cell cultures showed apoptotic or dead phenotype when 3μg/mL and 30μg/mL of anti-TNF-α, respectively, was included in anti-Ly-6A antibody treated YH16.33 cells (Figure 4B). This moderately reduced death in the cultures was not significant (p=0.6540) when compared to anti-Ly-6A cell cultures alone or in combination with the isotype control where an average of 72% and 73% of cells were apoptotic or dead (Figure 4B). Though there was a slight decrease in total apoptosis compared to anti-Ly-6A alone, treatment with anti-TNFα at both 3μg/mL (average 58%) and 30μg/mL (average 57%) concentrations did not decrease the populations of cells in apoptosis comparable to the “No Treatment” controls (approx. 72%).

**Figure 4.**
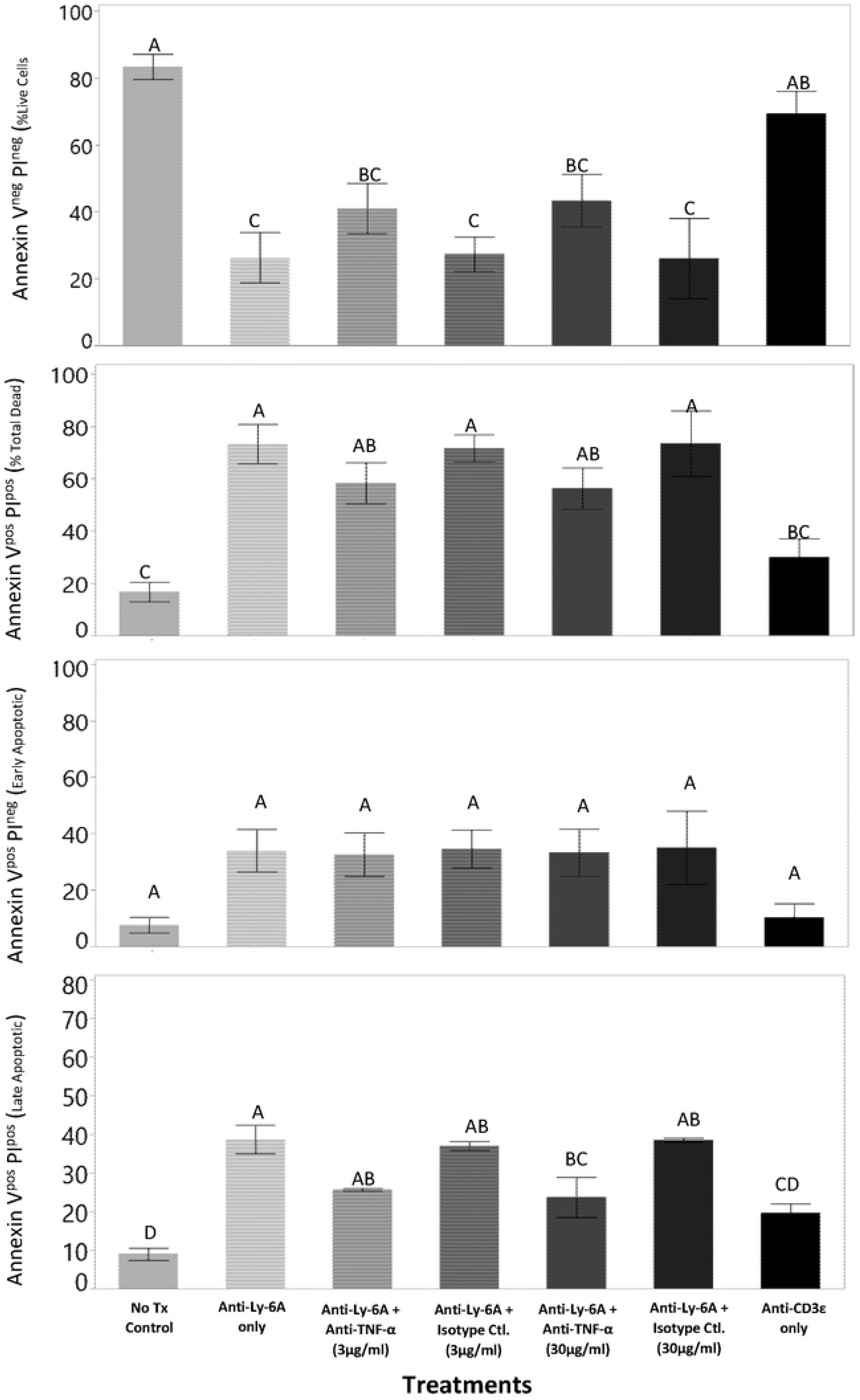
Effect of neutralizing anti-TNFα antibody on apoptotic cell death in YH16.33 in response to anti-Ly-6A stimulation. YH16.33 cells incubated with anti-Ly-6A, anti-CD3ε, and anti-TNFα or isotype control antibody or left untreated for 48 hours. Viable and apoptotic cells were analyzed after staining with Annexin V – FITC and Propidium Iodide followed by flow cytometric analysis. The figure represents data from 3 independent trials, with at least 5000 events per sample. The plotted mean of each treatment group shows percent viable (Annexin-V^neg^ PI^neg^), early apoptotic (Annexin V^pos^ PI^neg^) or late apoptotic/dead (Annexin V^pos^ PI^pos^), with error bars showing standard error. ANOVA analysis was performed with a post-test Tukey test. Groups with dissimilar connecting letters are significantly different from each other (p<0.05); n=3

An average of 34-35% of cells undergoing apoptosis (AnnexinV^pos^PI^neg^) were detected in cell cultures with anti-Ly-6A antibody, either alone or in combination with the isotype control antibodies at 3μg/mL and 30μg/mL concentration. This was significantly higher than what was observed in control untreated cell cultures (8%) and cells treated with anti-CD3ε (11%). Inclusion of neutralizing anti-TNFα antibodies at 3μg/mL and 30μg/mL (average 33%) did not alter the proportion of AnnexinV^pos^PI^neg^ cells in anti-Ly-6A stimulated cell cultures (average 33% at both anti-TNFα concentrations) when compared to the anti-Ly-6A stimulated cell cultures with isotype control antibodies (average 35%) (p>0.05) (Figure 4C). An average of 39% AnnexinV^pos^PI^pos^ cells were detected in cultures with anti-Ly-6A alone (Figure 4D). A similar percentage, 37% and 38%, of AnnexinV^pos^PI^pos^ cells was detected in anti-Ly-6A activated cultures with 3μg/mL or 30μg/mL isotype control antibodies, respectively. Isotype control antibody in cell cultures did not significantly alter anti-Ly-6A induced cell death. In contrast, inclusion of neutralizing anti-TNFα at 3μg/mL or 30μg/mL reduced the percentage of dead cells to 26% and 23%, respectively. This reduction was significant (p=0.0223) compared to the anti-Ly-6A treatment group (Figure 4D, this observation is consistent with the inclusion of isotype control antibody in anti-Ly-6A treated cells did not show significant change in percent apoptotic/dead cells compared to the controls (37-38%). The neutralizing TNFα reduces appearance of later apoptotic cells; however, the effects are not complete. Taken together, our data shows that engaging Ly-6A on CD4+ T cell line promotes secretion of TNFα, which in turn promotes apoptosis. These data suggest that Ly-6A triggered apoptotic cell death, in part, is triggered by an indirect, non-cell autonomous mechanism. In contrast, anti-CD3ε did not cause significant changes in the viability of YH16.33 cells (Figure 4A-D).

## DISCUSSION

Engaging Ly-6A, expressed on YH16.33 and other CD4^+^ T cell lines, with activating anti-Ly-6A antibody results in varied cellular responses ranging from growth inhibition, apoptosis and cytokine (IL-2 and IFN-γ) production [10]. Ly-6A signaling upregulates cyclin-dependent kinase inhibitor p27^kip1^ without altering expression of p53 [10]. Additionally, Ly-6A activates an intrinsic apoptotic cell death pathway by destabilizing mitochondria as evidenced by the release of cytochrome C in the cytoplasm and activation of caspases 3 and 9 [10]. The phenomenon of Ly-6A triggered growth inhibition, apoptosis and cytokine release by a clonal T cell line is paradoxical. In this study, we examined the relatedness of the three apparently distinct and conflicting functional responses. We hypothesized the role of non-cell autonomous mechanism in Ly-6A induced growth inhibition and apoptosis. We report that CD4^+^ T cell lines secrete TNF-α in response to Ly-6A stimulation, which in turn contributes to T cell growth inhibition and apoptotic cell death in the clonal T cell line. Additionally, TGF-β and GDF-10, a member of the TGF family, are minimally detected in these cell cultures and therefore less likely to contribute to the observed growth inhibitory and apoptotic functional responses.

CD4^+^ T cell lines, when stimulated through Ly-6A, are known to generate cytokines [13], including IFN-γ [24,25]. However, TNF-α production by clonal T cell lines after engaging Ly-6A proteins, and the subsequent biological effects, to our knowledge, are unknown. Our report adds TNF-α to a list of cytokines produced in response to Ly-6A activation. Importantly, anti-TNFα antibody blocking data (Figure 3) shows the contribution of TNFα in non-cell autonomous mechanism in Ly-6A-induced apoptosis/growth inhibition in CD4^+^ T cell line. These data are consistent with reports of TNFα, as a growth inhibitory and death factor [22] and in promoting induction of p27^kip1^ expression and caspase-3 activity [26]. Cellular effects of TNF-α on T cells, as described above, mirror Ly-6A signaling initiated upregulation of p27^kip^, release of cytochrome C in the cytoplasm and activation of Caspases 3 and 9 [10]. These findings contrast the observations where growth inhibition and apoptosis are not observed in the presence of anti-CD3 antibody despite the presence of TNFα and other inhibitory cytokines (TGF-β and GDF-10) (Figure 3; [10]). Future experiments are required to develop a mechanistic understanding of which TNF receptor (I or II) collaborates with Ly-6A in triggering growth inhibition/apoptosis and how engaging Ly-6A and the antigen receptor/CD3 may collaborate with different TNF receptors on the surface and/or access different signaling pathways that result in survival/no response or apoptosis/growth inhibition. It is intriguing that the two TNFα receptors, TNFRI and TNFRII have opposing biological activities [27] resulting in either apoptosis [28] or cell survival promoted by TNFα through the NFκB and the c-Jun pathways [29,30]. Incomplete reversal of growth inhibition/apoptosis suggest either incomplete neutralization of TNFα biological activity by anti-TNFα antibody and/or contribution by other cell death/signaling pathway, distinct from TNFα/TNFRI, in Ly-6A initiated growth inhibition/apoptosis. Our data taken together, suggest that the observed T cell growth inhibition and cell death is, in part, not only dependent on the presence of biologically active TNFα but also requires its combinatorial effect with continued Ly-6A signaling. As the presence of inhibitory cytokines in itself is not sufficient for growth inhibition, additional Ly-6A (not TCR/CD3) signaling is involved in the non-cell autonomous mechanism of T cell growth inhibition and apoptosis.

The Ly-6A protein is reported to interact with TGFβ receptors in mammary tumor cells and disrupt the GDF10 (a TGFβ-like cytokine)-dependent inhibitory signaling to enhance tumorigenicity in breast cancer cells [20]. However, we did not detect significant amounts GDF10 in T cell cultures stimulated with anti-Ly-6A antibody (Figure 1) suggesting its insignificant role in T cell growth inhibition. In addition, TGFβ, a known growth inhibitory cytokine, [31,32] was not abundantly detected in anti-Ly-6A-stimulated cell cultures (Figure 1) and therefore less likely to contribute to the observed growth inhibition/apoptosis. In contrast, significantly high free active and total TGFβ and GDF10 was detected in anti-TCRαβ/CD3 without detection of T cell growth inhibition. GDF10 and TGFβ are not produced by the transformed CD4^+^ T cell and therefore are less likely to play a biologically important role in Ly-6A driven growth inhibition of YH16.33 cells.

Ly-6A is a negative regulator of T cell expansion and poses as an immune checkpoint inhibitor involved in T cell clonal contraction phase on T cell response [33]. CD4^+^ T lymphocytes from Ly-6A-deficient mice show modest heightened proliferation rates compared to the controls [7]. Additionally, CD4^+^ T cells from Ly-6A transgenic mice displayed inhibited clonal expansion in response to a specific antigen [6]. The underlying mechanism of the growth inhibitory role of Ly-6A protein is unknown. Our data with the clonal T cell line proposes the role of TNFα and not the TGFβ and GDF10 in immune checkpoint inhibitory role of Ly-6A. Additional experiments will be required to experimentally examine this question.

## Acknowledgements

We thank Dr. Kenneth L. Rock for providing YH16.33 cell line and anti-Ly-6A/Sca-1 antibodies.

## Declarations

### Author Contributions

AP performed experiments, analyzed the data, performed the statistical analysis and partly wrote the manuscript. SM performed experiments and edited the manuscript. AKB pre-designed the project, coordinated the project, analyzed the data generated by AP and SM, wrote the manuscript and edited the manuscript. All authors have read and approved the final manuscript.

### Ethical approvals and consent to participate

Not applicable

### Authors information

Correspondence address: Anil K Bamezai, Department of Biology, Villanova University, 800 E Lancaster Avenue, Villanova, PA 19085, Phone No. 610.519.4847; Fax No: 610.5197863

First author: Akshay Patel

Current address: MD/PhD Program, Department of Biochemistry & Molecular Biology, SUNY Upstate Medical University, Syracuse, NY 13210

E.mail

Co-author: Sarah Moxham

Current address: Department of Classics, 800E Lancaster Avenue, Villanova University, Villanova, PA

E.mail

## Abbreviations

Ly-6: Lymphocyte antigen-6
CD3: Cluster differentiation antigen 3
CD4: Cluster differentiation antigen 4
TCRαβ: T cell receptor-alpha beta
TNF-α: Tumor Necrosis Factor-α
TGF-β: Transforming Growth Factor-β
GDF-10: Growth Differentiation Factor 10
μM: micromolar
mM: millimolar
IACUC: Institution animal care and use committee
RPMI 1640: Roswell Park Memorial Institute *1640* Medium
HEPES: (4-(2-hydroxyethyl)-1-piperazineethanesulfonic acid)
FL 1-4: fluorescence channel 1-4
MTT: 3-(4,5-dimethylthiazol-2-yl)-2,5-diphenyl tetrazolium bromide
ELISA: Enzyme Linked Immunosorbent Assay
ANOVA: analyses of variance

